# Saccade vigor partly reflects the subjective economic value of the visual stimulus

**DOI:** 10.1101/770560

**Authors:** Tehrim Yoon, Afareen Jaleel, Alaa A. Ahmed, Reza Shadmehr

## Abstract

Decisions are made based on the subjective value that the brain assigns to options. However, subjective value is a mathematical construct that cannot be measured directly, but rather inferred from choices. Recent results have demonstrated that reaction time and velocity of movements are modulated by reward, raising the possibility that there is a link between how the brain evaluates an option, and how it controls movements toward that option. Here, we asked people to choose among risky options represented by abstract stimuli, some associated with gain, others with loss. From their choices in decision trials we estimated the subjective value that they assigned to each stimulus. In probe trials, they were presented with a single stimulus at center and made a saccade to a peripheral location. We found that the reaction time and peak velocity of that saccade varied roughly linearly from loss to gain with the subjective value of the stimulus. Naturally, participants differed in how much they valued a given stimulus. Remarkably, those who valued a stimulus more, as evidenced by their choices in decision trials, tended to move with greater vigor in response to that stimulus in probe trials. Thus, saccade vigor partly reflected the subjective value that the brain assigned the stimulus. However, the influence of subjective value on vigor was only a modest predictor of preference: vigor in probe trials allowed us to predict choice in decision trials with roughly 60% accuracy.

**New and Noteworthy:** We found that saccade vigor tends to vary monotonically with subjective value: smallest for stimuli that predict a loss, and highest for stimuli that predict a gain. Notably, between-subject differences in valuation could be gleaned from the between-subject differences in their patterns of vigor. However, the influence of subjective value on vigor was modest, allowing partial ability to infer subjective value for the purpose of predicting choice in decision trials.

---

> “A true theory of economy can only be attained by going back to the great springs of human action --- the feelings of pleasure and pain.”
>
> — William Stanley Jevons (1866)

## Introduction

Theory of subjective value was introduced in the 19^th^ century to account for the fact that in voluntary transactions, each party values the goods, labor, or money that they receive more than the goods, labor, or money that they provide (Jevons, 1866;Menger, 1871). The theory posited that subjective value is not specified by an objective property of the good, but rather the incremental increase in pleasure that an individual assigns to acquisition of that good (Jevons, 1866). Although subjective valuation is an important aspect of behavioral economics, it is an abstract quantity that cannot be measured directly. Rather, it must be inferred from decisions that individuals make (von Neumann and Morgenstern, 1944), often in scenarios involving lotteries and risky options.

A serendipitous discovery in motor neuroscience has been the observation that factors that affect preference, such as reward and effort, also affect movements (Shadmehr et al., 2019). For example, in goal-directed movements, people and other primates move with a shorter reaction time and greater velocity towards stimuli that they associate with greater gain (Kawagoe et al., 1998;Milstein and Dorris, 2007;Xu-Wilson et al., 2009;Yoon et al., 2018;Summerside et al., 2018). Recent work (Sedaghat-Nejad et al., 2019) has shown that if presentation of a stimulus results in a reward prediction error, the movement that ensues tends to be expressed with greater vigor (defined as the reciprocal of reaction-time plus movement-duration). Intriguingly, reward prediction error is the principal variable that modulates dopamine release (Schultz et al., 1997;Bayer and Glimcher, 2005), and stimulation of dopamine around movement onset tends to increase vigor (da Silva et al., 2018). Thus, both the process of learning subjective value from reward prediction error, and control of movement vigor, depend on dopamine, raising the intriguing possibility that the vigor with which an individual moves toward an option is partly influenced by the subjective value that they assign to that option.

Previous work has established that when people are presented with a decision between two options, their deliberation time is a measure of their strength of preference: participants typically decide sooner if they prefer one stimulus much more than another (Spiliopoulos and Ortmann, 2018;Konovalov and Krajbich, 2019). Thus, these works have demonstrated that certain aspects of behavior during decision making are related to the difference in the subjective value of the two options.

Here, we asked a different question: suppose one could only observe movements during presentation of a single stimulus, but not during decision-making. Can one infer subjective value from the movement vigor toward single stimuli A and B, and then predict choice when the subject decides between A and B? If so, how well might movement vigor in single stimulus trials allow one to predict subjective value, and thus choice in decision trials?

It is possible that vigor may not reflect subjective value, but rather an aspect of attention allocated to the stimulus. For example, both the stimulus that promises a gain and the stimulus that foretells a penalty are important and will garner more attention than stimuli that promise smaller gain and loss. In this scenario, vigor will not increase monotonically with subjective value, but rather produce a U-shaped function, becoming large for both gains and losses.

There are plausible neural mechanisms that support this alternate hypothesis. Saccade vigor is partly modulated by the excitatory inputs that the superior colliculus receives from the cortical regions which compute subjective value: the frontal eye field (FEF) (Hanes and Schall, 1996;Heitz and Schall, 2012;Glaser et al., 2016) and the lateral intraparietal area (LIP) (Platt and Glimcher, 1999;Louie and Glimcher, 2010). LIP neurons that encode stimulus value exhibit greater activity both when the stimulus promises a large reward, and when the stimulus promises a large penalty (Leathers and Olson, 2012). Some of these neurons exhibit sensitivity to both novelty and value (Foley et al., 2014). Furthermore, some dopamine neurons increase their activity when the stimulus promises reward, whereas others increase their activity for both punishment and reward (Matsumoto and Hikosaka, 2009). Thus, the neural activity that could modulate saccade vigor shows positive sensitivity to gain, as well as loss. This leads us to the question of whether vigor monotonically reflects valuation over a range that includes both losses and gains, or is vigor a U-shaped function of value.

Here measured saccades in a task where humans learned to associate value to 10 abstract stimuli, each paired with a different magnitude of loss or gain. By design, the task involved learning, which we hoped would result in some participants valuing a given stimulus highly, whereas others would value it less. In probe trials, we presented one stimulus at random and measured saccade vigor. In decision trials, the participants deliberated between various stimuli and made a choice, from which we also inferred the subjective value that they assigned to each stimulus. We found that in probe trials, saccade vigor was lowest for stimuli that promised a loss, and highest for stimuli that promised a gain. However, while vigor was clearly affected by subjective value, this effect was modest: using vigor in probe trials as a proxy for subjective value, we could correctly predict about 60% of the choices made in decision trials.

## Materials and Method

Healthy participants (n=24, 26.3±8.2 years old, mean±SD, 8 females) with no known neurological disorders and normal color vision sat in a well-lit room in front of an LED monitor (59.7 × 33.6 cm, 2560 × 1440 pixels, light gray background, frame rate 144 Hz) placed at a distance of 35 cm. Their head was restrained using a bite bar. They viewed visual stimuli on the screen, and we measured their eye movements using an EyeLink 1000 (SR Research) infrared recording system (sampling rate 1 KHz). Only the right eye was tracked. All participants were naïve to the paradigm. The experiments were approved by the Johns Hopkins University School of Medicine Institutional Review Board, and all participants signed the written consent form approved by the board. Participants were paid $15/hour regardless of any behavioral outcome. One participant was excluded from the results presented here because their performance in the task was at chance level, suggesting that they did not learn to assign value to the various stimuli.

### Stimulus properties

We performed an experiment in which through experience, people learned the value of 10 abstract visual stimuli. Each stimulus was a 2° × 2° colored square, designated with a “+” or “-“ (Fig. 1B). Each square was randomly assigned to a point distribution, with a mean that ranged from loss of 5 points to gain of 5 points. The points associated with each color were selected randomly on each trial from a beta distribution with parameters α = β = 2, scaled so that each color was associated with a single mean: −5, 4, …, +5. The plus and minus indicator at the center of the square noted the sign of the mean of the distribution. The color-to-point relationship was selected randomly for each subject, but remained consistent throughout the experiment. For example, the plus yellow square in Fig. 1B was associated with a distribution with mean equal to gain of 4 points, and the minus yellow square was associated with mean equal to loss of 4 points. In addition to these 10 colored squares, a black square with “0” at the center was associated with exactly 0 points. Thus, the experiment design used abstract stimuli that through experience, participants learned to associate with points. We hoped that this would produce a wide diversity in subjective values that the participants assigned to a given stimulus, allowing us to test whether movement vigor was a predictor of the between participant differences in subjective value.

**Fig. 1.**
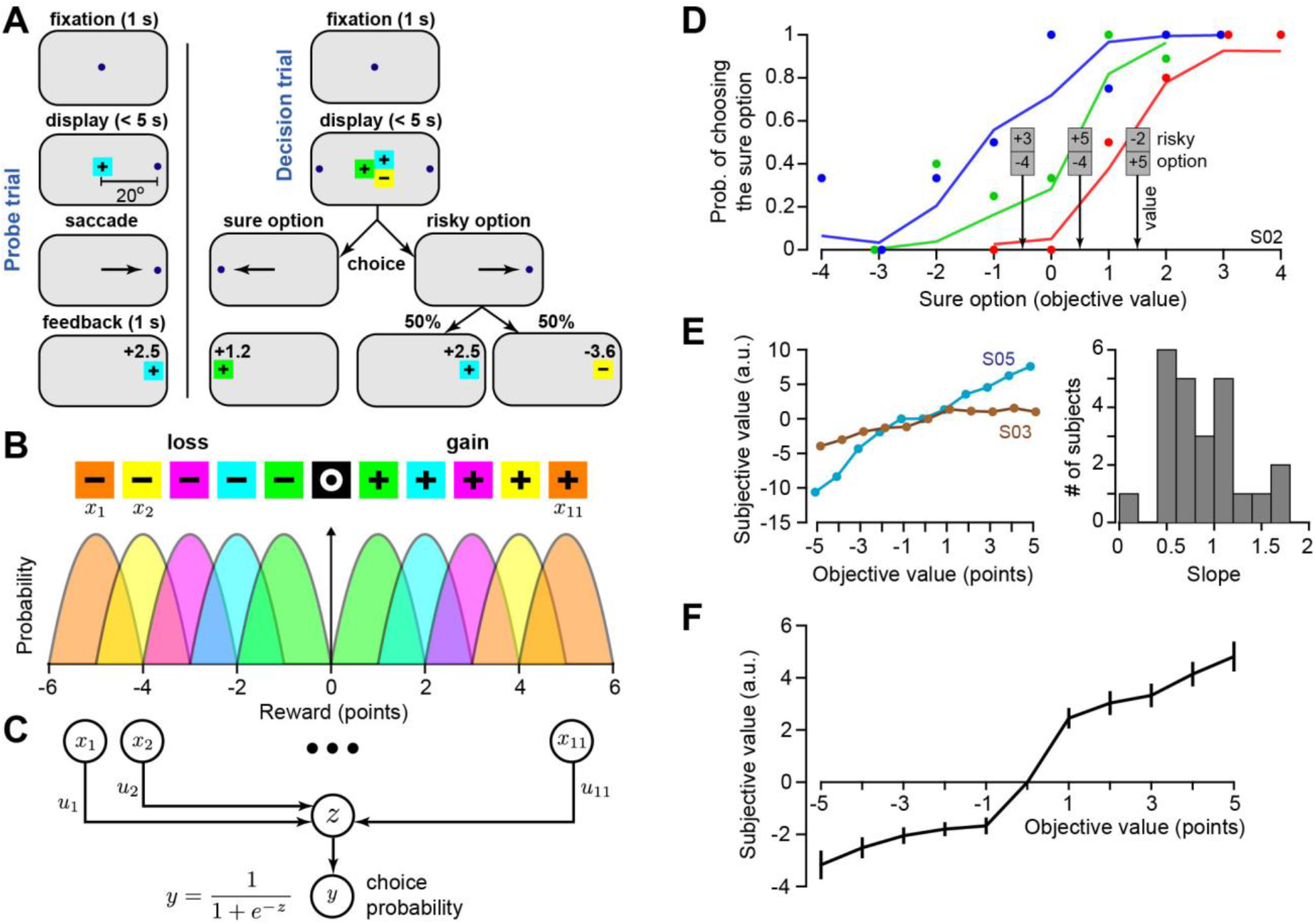
Estimating subjective value of abstract stimuli. **A**. In probe trials, a single stimulus was presented at center and a dot that served as saccade target ±20°. By making a saccade, the participants earned the points associated with that stimulus (gain or loss). In decision trials, a single stimulus representing a sure bet, and two stimuli representing a risky bet, appeared at center. The participants made a choice by making a saccade to one side or the other. **B**. The stimuli consisted of 11 boxes. The colored stimuli were associated with gain or loss (indicated with the plus or minus), each with a distribution as shown. The black stimulus was always associated with zero points. **C**. We used a neural network to model the decision-making process. The input **x** was an 11 element vector, with each element representing one of the stimuli *x*_1_, ⋯, *x*_11_ starting from the most negative to the most positive, and the black square (0 points) being the sixth element. On each trial, the input vector **x** was set so that one element had value of −1 for the sure stimulus, two elements had value of +0.5 for the pair of risky stimuli, and 0 for the remaining elements. The weight vector **u** represented the subjective value of each stimulus. Variable *z* was determined by Eq. (1), and the output of the network was the probability of picking the sure option. **D**. Some of the choices made by a participant. The colored dots indicate the stimuli that were presented for the risky option, the x-axis is the point value of the sure option, and the y-axis is the probability of picking the sure option by this participant. For example, the green dots show the probability that the participant picked the sure option when the risky option was +5 and −4 stimuli. The expected value of risky option was 0.5. This participant tended to pick the sure option if that option had a value greater than 0.5. **E**. There was diversity in the subjective valuation that the participants learned to assign to the stimuli. The left panel shows subjective values that two participants learned. The right panel shows the distribution of the slope of the subjective vs. objective values across the participants. **F**. Subjective values across all participants. Error bars are SEM.

**Fig. 2.**
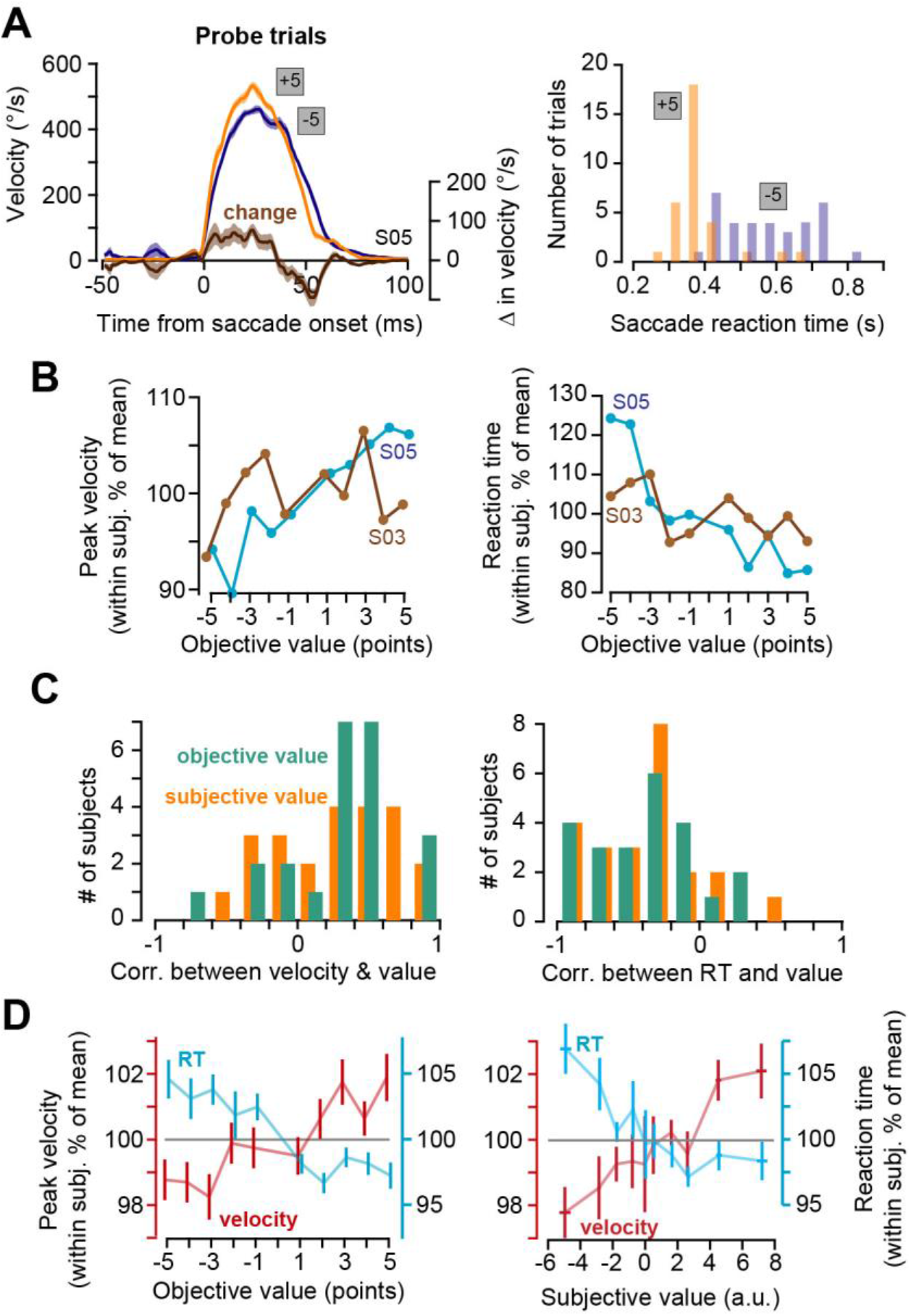
Saccade characteristics in probe trials. **A**. Data from a single participant. Left subplot shows saccade velocity in probe trials in which the stimuli were +5 and −5. Right subplot shows saccade reaction times for the same two stimuli. Error bars are within subject SEM. **B**. Peak velocity and reaction time in probe trials as a function of objective stimulus value for two participants (same individuals as those shown in Fig. 1E). Saccade vigor in S05 appeared to be more strongly modulated by stimulus value than S03. **C**. The distribution of correlation coefficients between velocity and stimulus value, and between reaction time and stimulus value. **D**. Peak saccade velocity and reaction times as a function of objective and subject values. Saccade data are normalized within subject with respect to their own mean in probe trials. Error bars are between-subject SEM.

### Decision and probe trials

The experiment contained two types of trials, randomly intermixed. Both types of trials (Fig. 1A) began with a center fixation period that lasted for 1 sec and ended with a beep (1 KHz). In decision trials, the fixation point was replaced with three different colored stimuli. One stimulus appeared alone and represented a sure bet (100% probability of acquiring the points associated with that stimulus). The other two stimuli appeared together and represented a risky bet (each with 50% probability). The participant had 5 seconds to indicate their choice by making a saccade in direction of their choice to targets (square, 0.5 × 0.5°) that appeared on the horizontal axis at ±20°. Once the saccade concluded, the stimuli at center were erased and the trial consequences were displayed for 1 sec: the earned stimulus was displayed at the dot location along with text that indicated the number of points acquired. The points were drawn from the random distribution associated with the colored stimulus. Failure to make a choice within the time limit resulted in loss of 10 points. The trial ended with the display of the color stimulus and the amount of points gained or lost for that trial (duration of 1 sec).

In probe trials, the fixation point was removed, a single stimulus (chosen at random from the 10 colored stimuli) was displayed at center, and a dot appeared on the horizontal axis (at ±20°). This was the instruction for the subject to make a saccade to the dot. Once the saccade concluded, the stimulus at center was erased and displayed at the dot location, along with text that indicated the number of points that the subject had gained or lost for the trial. As in the decision trials, the points were drawn from the random distribution associated with the colored stimulus. Thus, trials included stimuli that were associated with gain or loss, and by making a saccade, the subject earned that gain or loss. Failure to make a saccade resulted in loss of 10 points.

### Experiment design

Before the start of the experiment, the participants were instructed that there were 10 stimuli consisting of two sets of 5 colored boxes that represented points that could be gained or lost on each trial. “Each color will indicate how many points you will gain or lose. Black box will always give zero points when chosen. Boxes with plus signs will add to your score, while boxes with minus signs will decrease your score. For example, if orange box with plus sign indicates gain of 10, orange box with minus sign will indicate loss of 10.”

The experiment consisted of 11 blocks, each with 100 trials. The first block was a training block and began with 100 points and included only probe trials. This first block served to teach the participants the points associated with the various stimuli. The remaining 10 blocks each had 40 probe trials and 60 decision trials, distributed randomly. The total score was reset to 100 at the start of the 2^nd^ block. For probe trials, each of the 10 colored squares was presented with equal frequency within each block, distributed randomly. In probe trials, the direction of the dot was chosen randomly at left or right but with equal frequency within each block.

In a decision trial, we randomly picked 3 stimuli from among the 11 stimuli. We presented the medium valued stimulus as the sure bet and the other two stimuli (one loss, and the other gain) as the risky bet. Participants were not provided any information about the value of the stimuli and thus had to make their decisions solely based on consequences of previous trials. The side that represented the sure bet was random and chosen with equal left-right frequency for each block. Following completion of the second block, the final score of the previous block was carried over as the starting score of the next block. At the conclusion of every 4^th^ trial, the total score earned was displayed at center fixation.

### Data analysis

Eye position data were filtered with a second-order Savitzky-Golay filter (frame size 11, degree 3). Saccade onset and offset were determined in real time with 20°/s threshold. We identified valid saccades as those that occurred between stimuli with start and endpoints that were within 5° of the boundaries of the start and end images (to account for the fact that participants were not specifically instructed to fixate on a precise location). For probe trials, we excluded reaction times that were larger than 1 sec.

Our objective was to test whether behavior in probe trials reflected the subjective value that we had estimated from decision trials. Thus, we analyzed vigor of saccades only in probe trials, and inferred subjective value based on choices made in decision trials. Statistical testing relied on linear mixed-effect models. In each model, the dependent variables were saccade peak velocity and reaction time, fixed effects were stimulus objective value and subjective value, and random effects were individuals. Dependent variables were normalized for each individual by dividing the measured value by the within subject mean. Statistics were performed on normalized dependent variables.

### Estimating subjective value of stimuli

The objective value of each stimulus was set by the mean of the point distribution associated with each colored square (Fig. 1B). The participants formed subjective values, and we inferred these values based on the choices that they made in decision trials.

In a decision trial, the choice was between a sure option (a single stimulus) and a risky option (two stimuli, 50% chance of each). To model the choices that participants made, we designed a one-layer perceptron network that had as its input the three stimuli that were available on each trial. The output of the network was the probability of picking the sure option (Fig. 1C). The input **x** was an 11 element vector, with each element representing one of the stimuli *x*_1_, ⋯, *x*_11_ starting from the most negative to the most positive, and the black square (0 points) being the sixth element. On each trial, the input vector **x** was set so that one element had value of −1 for the sure stimulus, two elements had value of 0.5 for the pair of risky stimuli, and 0 for the remaining elements. The weight vector **u** represented the subjective value of each stimulus, and was also an 11 element vector. A linear combination of the available stimuli were represented with variable *z*:

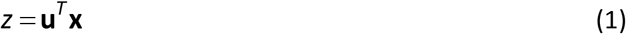

For example, if in a given trial the sure option was stimulus *x*_4_, and the risky option was stimuli *x*_2_ and *x_7_*, then *z* = 0.5(*x*_2_ + *x*_7_) – *x*_4_. In other words, the variable *z* represented the difference between the subjective values of the two options. This was then transformed via a logistic function that produced an output *y* that represented the probability of picking the sure option:

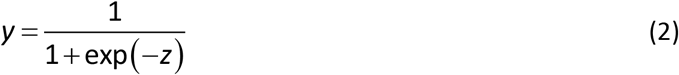

Our objective was to estimate the subjective value that the participant had assigned to each stimulus, represented via the weight vector **u**. We assumed that the subjective value of the zero stimulus (the sixth element of **u**) was exactly zero. To find the remaining weights, we used a binary cross-entropy loss function:

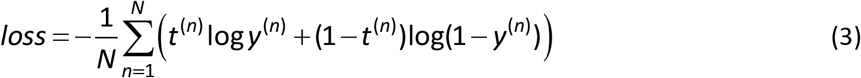

In the above equation, *N* is the total number of decision trials (600). Binary variable *t*^(*n*)^ represented the actual decision of the participant on trial *n*: *t*^(*n*)^ = 1 for choosing the risky option, and *t*^(*n*)^ = 0 for the sure option. To find **u**, we differentiated Eq. (3) with respect to **u**, thus providing a stochastic gradient descent estimate of the subjective values. We stopped the algorithm when the norm of change of the subjective value Δ**u** for was less than 10^−4^.

### Using vigor in probe trials to predict choice in decision trials

Once we determined the reaction time and peak saccade velocity associated with a given stimulus in the probe trials, we asked whether vigor could serve as a proxy for subjective value. To evaluate the accuracy of such a policy we used two different approaches: a winner take all approach that predicted choice in the decision trials, and a likelihood estimate approach that predicted probability of choice in decision trials.

In the winner take all approach, for each stimulus in the probe trials we computed saccade velocity and reaction time, imagined that subjective value of the stimulus was set by these variables, and then on each decision trial used these measures to predict choice. For example, to evaluate the vigor policy, we assigned subjective value to the 11 stimuli based on vigor on the probe trials, and then used this to predict choice of the participant in each of the decision trials: pick the option that has the larger vigor estimated subject value (100% of the vigor for the sure option stimulus, vs. sum of 50% of vigor for each of the risky option stimuli). We compared the accuracy of this vigor-based policy with a policy that made choices based on subjective values that were estimated based on the actual decisions of each subject. To predict outcomes, we used the actual options faced by each participant. We used Wilcoxon signed-rank test to compare performance of the various policies.

In the likelihood estimate approach, we began by setting the vector **u** to be equal to the mean reaction time (or peak velocity) for the various stimuli in the probe trials. We then used this vigor based estimate of subjective value to predict the probability that the participant would pick the sure option in a given decision trial:

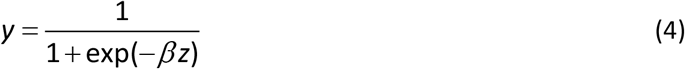

In the above formulation, the term *β* appears because unlike Eq. (2), *z* in Eq. (4) has units that are different than subjective value. We used Eq. (3) to guide the gradient descent procedure for finding *β* for each participant. Thus, using vigor as an estimate for subjective value, we used Eq. (4) to predict the probability that on a given trial the participant would pick the sure option. We then evaluated the goodness of this policy by computing the negative log likelihood (NLL). We compared the vigor based policy to a random policy. NLL for the random policy was one in which the probability of choosing the sure option was 0.5 (equivalent to having elements of **u** equal to each other).

## Results

The participants were presented with colored stimuli that were probabilistically associated with gain or loss (Fig. 1B). In the baseline set of probe trials, they were repeatedly exposed to the color-point pairings, and then in a subsequent set of decision trials they expressed their preferences.

On decision trials, participants were presented with a sure option and a risky option (Fig. 1A, right column). Some of the choices made by a representative participant are shown in Fig. 1D. The x-axis of this plot presents the objective value of the sure option on various trials (mean of the point distribution associated with that stimulus, Fig. 1B). The y-axis of Fig. 1D presents the probability that the participant selected the sure option. The various colored dots indicate the stimuli that were presented for the risky option. For example, the green dots show the probability that the participant picked the sure option when the risky option consisted of the colors associated with +5 and −4 points. The expected value of the risky option was 0.5. Indeed, this participant tended to pick the sure option if that option had a value greater than 0.5. In another example, the red dots show the probability that the participant picked the sure option when the risky option was −2 and +5 points. In this case, the risky option had an expected value of 1.5. The participant now picked the sure option when that option had a value that was greater than 1.5.

We used a one-layer neural network to model the choices that each participant made in the decision trials and inferred the subjective value that they had assigned to each stimulus (Eq. 2). On each trial, given the sure and two risky stimuli, the network predicted the probability that the subject would pick the sure option (lines in Fig. 1D). We trained the network with the actual choices that the subject had made. The loss function (Eq. 3) was guaranteed to minimize the difference between choices observed and choices predicted. After fitting the network to the data of each subject, the weight vector **u** provided the estimate of the participant’s subjective value for each stimulus.

Examples of subjective values inferred from choices made by two participants are illustrated in Fig. 1E (left panel). Participant S03 learned a shallow function, distinguishing between positive and negative valued stimuli, but not distinguishing well within stimuli that were positive or negative. In contrast, participant S05 learned a steep function that distinguished well within negative and positive stimuli. Thus, among the participants there was diversity in the value that they assigned each stimulus, as shown by the distribution of subjective vs. objective value slope (0.88±0.40, mean±SD, Fig. 1E, right panel). As we will see, this diversity played an important role in the question of whether vigor was driven by subjective value.

The average pattern of subjective value is displayed across the participants in Fig. 1F. We found that on average, subjective value strongly correlated with objective value of the stimuli (r^2^ = 0.72; p<10^−30^). Within participant analysis of subjective value revealed a main effect of objective value (F(1,262)=585.7, p<10^−30^), demonstrating that as the objective value of the stimuli increased, so did the subjective value that the participants had assigned to them. Thus, the participants learned the task.

These data indicated that the participants generally learned to assign value to the abstract stimuli, resulting in subjective values that increased with objective values. However, there were also differences among participants, with some learning steep value functions, while others learned shallow functions.

### Vigor increased with stimulus value

In probe trials a single colored stimulus appeared at center, indicating the value of that trial, and simultaneously a dot appeared on the periphery, indicating the saccade target. Presentation of the colored stimulus served as the go cue. The dependent variables were reaction time and velocity of the ensuing saccade. Participants had 5 seconds to make a saccade to the peripheral dot and by doing so earned the loss or gain that was associated with the stimulus. Notably, the loss that was indicated by the negative valued stimuli was always less than the large penalty (10 points) that would be applied if the participants did not make the correct saccade.

The left subplot of Fig. 2A illustrates saccade velocities for one participant in probe trials for +5 and −5 stimuli. The reaction times for these two stimuli are presented in the right subplot of Fig. 2A. In response to the higher valued stimulus, this participant produced a saccade that had a shorter reaction time, and a higher peak velocity.

To examine these trends across the participants, we normalized saccade peak speed and reaction times for each individual with respect to their own mean as measured across probe trials (Haith et al., 2012). Data from two participants are presented in Fig. 2B. Participant S05 exhibited peak velocity and reaction time that correlated strongly with stimulus objective value (velocity: r=+0.93, p<10^−4^; RT: r=−0.96, p<10^−5^). In contrast, participant S03 exhibited saccade vigor that was poorly correlated with objective value (velocity: r=+0.39, p=0.26; RT: r=−0.48, p=0.16). The distribution of correlation coefficients for all participants is plotted in Fig. 2C. The correlation coefficients between reaction time and subjective value had a mean of −0.38±0.076 (Wilcoxon signed-rank test, p=2.1×10^−4^). With respect to objective value, this distribution had a mean of −0.42±0.072 (Wilcoxon signed-rank test, p=3.2×10^−4^). The distribution of correlation coefficients between peak velocity and subjective value had a mean of 0.29±0.082 (Wilcoxon signed-rank test, p=0.0051). With respect to objective value, this distribution had a mean of 0.28±0.085 (Wilcoxon signed-rank test, p=0.0032). To examine these results together, we binned the vigor data based on the objective value of the stimulus (10 bins, one per stimulus, Fig. 2D) and found that saccade velocity increased with objective value of the stimulus (within subject effect, F(1,228)=28.6, p=2.2×10^−7^). Similarly, reaction time decreased with objective value of the stimulus (within subject effect, F(1,228)=50.6, p=1.4×10^−11^). Thus, although there was diversity among the participants, saccade vigor in probe trials was significantly affected by objective value of the stimulus.

We next tested the effects of subjective value on vigor. We observed that as subjective value increased, saccade velocities increased (within subject effect, F(1,228)=33.6, p=2.3×10^−8^), and reaction times decreased (within subject effect, F(1,228)=62.2, p=1.3×10^−13^), as shown in the right part of Fig. 2D. [To make this plot, we began with the distribution of subjective values across all participants, and then sampled that distribution into 10 bins of equal probability. Thus, the bins have error bars in both x- and y-dimensions.] Together, these data demonstrated that vigor in probe trials was not a U-shaped function of stimulus value. Rather, vigor tended to be smallest for stimuli that were associated with loss, and largest for stimuli that were associated with gain.

Given the between-subject diversity in the relationship between vigor and stimulus value in probe trials (Fig. 2C), we wondered whether there was some characteristic of participants in decision trials that dissociated their vigor modulation in probe trials. One clue was that some participants learned a steep value function, while others learned a shallow function (Fig. 1E). Indeed, we found that the slope of subjective to objective values was modestly correlated with the slope of saccade velocity with respect to subjective values (slope of velocity vs. subjective value compared to slope of subjective value vs. objective value, r=0.49, p=0.019). That is, the participants whose saccade vigor was more strongly modulated by stimulus value in probe trials tended to have learned a steeper value function, as inferred from their choices in decision trials.

In summary, we observed that in probe trials saccades had reaction times that decreased with subjective value, and peak velocities that increased with subjective value. However, there was diversity in the strengths of these relationships. It appeared that saccade vigor was more strongly modulated by stimulus value in those participants who had also learned a steeper value function.

### Between subject differences in subjective value influence between subject differences in vigor

Some participants learned to assign a large subjective value to a stimulus, while others assigned a lower value to the same stimulus. Could this between-subject difference in valuation be gleaned from the vigor patterns?

To examine this question, we described our hypothesis via a graphical model (Fig. 3A). In this model, choice depended on subjective value, which in turn depended (through learning) on the objective value of the stimulus. In our null hypothesis (H0, Fig. 3A), the objective value affected vigor, whereas subjective value affected choice. In our main hypothesis (H1, Fig. 3A), objective value affected subjective value, which in turn affected both choice and vigor. Under H1, if a subject had learned to associate a small subjective value with a stimulus, then their vigor would be low in response to that stimulus. However, if that same stimulus was valued highly by another subject, then their vigor would be high. Thus, to test this hypothesis, we kept objective value constant and asked whether changes in subjective value across participants modulated saccade vigor.

**Fig. 3.**
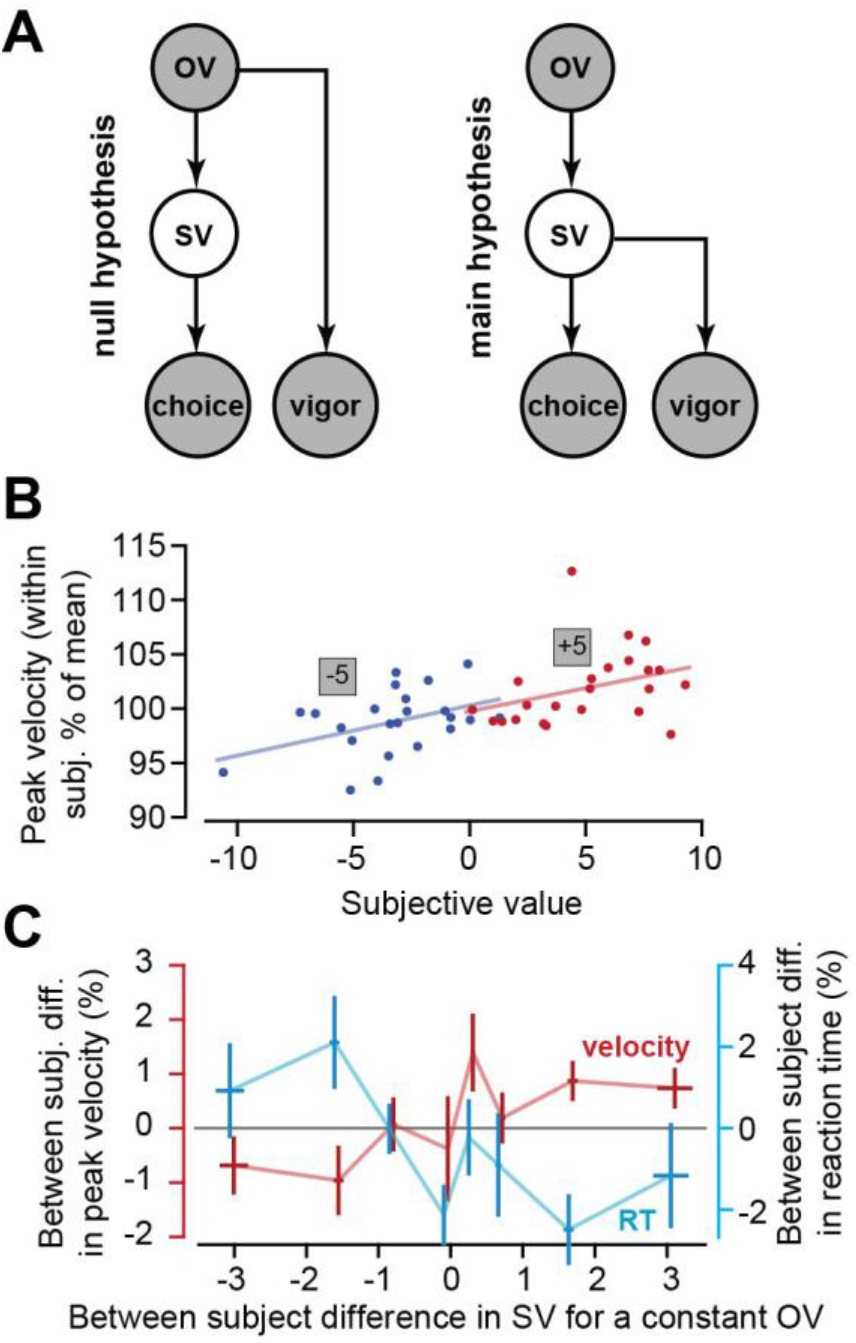
Statistical relationship between vigor, objective value (OV), and subjective value (SV). **A**. Graphical model representing two hypotheses. Each circle is a random variable. OV is objective value, and SV is subjective value. The filled circles are measured variables. The unfilled circle is not measured but estimated. In the null hypothesis, choice depends on SV, and both SV and vigor depend on OV. In the main hypothesis, SV affects both choice and vigor. **B**. To evaluate merits of the hypotheses, we kept OV constant and measured variability in vigor as a function of variability in SV. In this plot blue dots are peak saccade velocity as a function of subjective value (each dot is a participant), for the fixed objective value of −5. The red dots are for the fixed objective value of +5. For a given stimulus, some participants assigned a high SV, while others assigned a low SV. Vigor appeared to be higher when the participant assigned a high SV to the stimulus. **C**. To consider the data in part B across stimuli, we found the mean of the dot distribution for each stimulus (for example, the mean for the stimulus OV=+5), and then represented each dot with respect to the within stimulus mean. The result revealed that for a constant OV, a change in SV produced a change in vigor, thus rejecting the null hypothesis.

To help explain how we tested this hypothesis, Fig. 3B illustrates peak velocity in probe trials as a function of subjective value for two different stimuli. For the +5 stimulus, some participants assigned a large value, while others assigned a small value. Similarly, for the −5 stimulus, there was diversity in assignment of subjective values. However, individuals that assigned larger subjective value to a given stimulus also appeared to move with greater velocity in response to that stimulus (similar positive slopes of the red and blue lines in Fig. 3B).

To test for the consistency of this relationship, for each stimulus (constant objective value) we measured the vigor of a participant (i.e., the y-value of a point in Fig. 3B with respect to the mean of the points with the same color), and the subjective value that they had assigned (i.e., the x-value of a point in Fig. 3B with respect to the mean of the points with the same color). Thus, given a constant objective value, we measured how the between-subject differences in subjective value affected between-subject differences in vigor (Fig. 3C). We found that given a constant objective value, an increase in subjective value produced a reduction in reaction time (F(1,228)=8.7, p=0.0036), and an increase in peak velocity (F(1,228)=8.1, p=0.0047).

These results suggested that between-subject differences in valuation of a stimulus in decision trials could be partially inferred from the between-subject differences in saccade vigor in decision trials: participants who learned to associate a greater value to a given stimulus also tended to exhibit a greater modulation of vigor in response to that stimulus.

### Vigor was a modest predictor of choice

We next asked how well vigor measurements in probe trials could be used to predict choices that individuals made in decision trials. To predict choice, we used only the vigor data in probe trials. The vigor measurements produced two policies: a policy that assigned subjective value based on reaction time in probe trials, and another policy that assigned subjective value based on peak velocity in the same trials. For example, given the probe trial data for a participant, the reaction time policy assigned a subjective value to the various stimuli, which we then used to predict choice in decision trials for that participant. This served as the winner take all approach. In addition, we considered a likelihood approach in which we predicted the probability that the participant would pick the sure option based on their vigor patterns in the probe trials.

For the winner take all approach, we divided the decision trials into easy and hard based on the difference in the objective value of the sure and risky options: easy trials were denoted by objective value difference of 1 point or more, and hard trials were denoted by objective value difference of less than 1 point. We quantified accuracy of the vigor policies based on the number of correct predictions that the reaction time and the velocity policies made. To define an upper bound on prediction accuracy, we also quantified performance of a policy that relied on the neural network that had fit the actual choices (termed logistic fit).

The results of the policy comparisons are shown in Fig. 4A. We found that for hard choices, a velocity based policy performed no better than chance (Fig. 4A, Wilcoxon signed rank test, p=0.99). However, for the same hard choices a reaction time policy performed significantly better than chance (Wilcoxon signed rank test, p=0.0042). For the easier choices, both the velocity based policy and the reaction time policy performed significantly better than chance (Fig. 4A, left subplot, velocity Wilcoxon signed rank test p=0.0225; reaction time Wilcoxon signed rank test p=5.5×10^−4^).

**Fig. 4.**
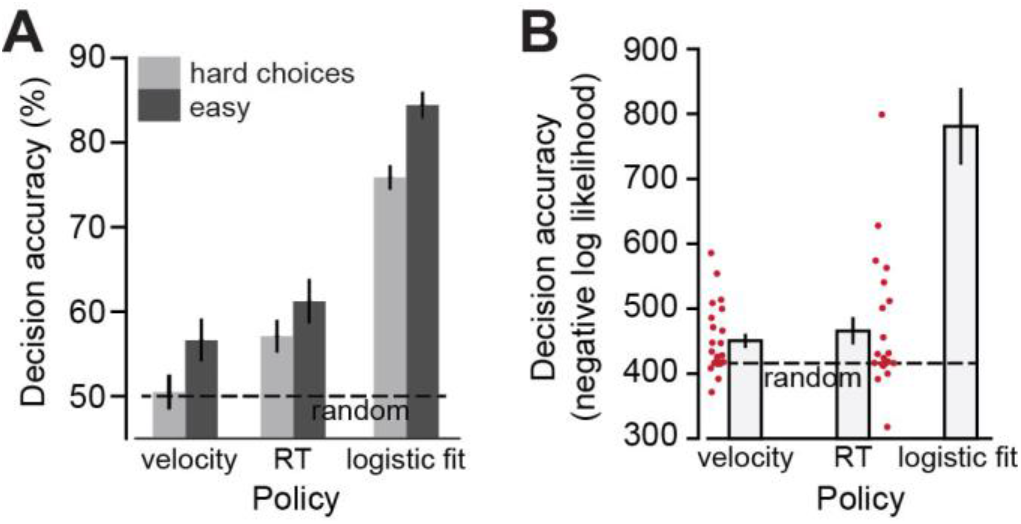
The ability to predict choice in decision trials from vigor patterns in probe trials. **A**. Hard choices are those in which the difference between the objective values of the two options was 1 point or less. Easy choices are those in which this difference was greater than 1 point. The velocity and reaction time policies computed subjective values based on probe trials, and then predicted choices in decision trials. Logistic fit policy fit the data in decision trials and predicted choice in the same data set, thus representing the ceiling for a model based prediction. B. Decision accuracy as measured via a log-likelihood estimate. From the vigor data in probe trials we predicted the probability that the participant would pick the sure option in a given decision trial (Eq. 4), and then evaluated the goodness of the policy by computing the negative log-likelihood. The dashed line indicates performance of a random policy.

In addition to predicting choice via winner take all, we also used vigor based estimates of subjective value to compute the probability of choosing the sure option (Eq. 4). We estimated the goodness of the vigor based predictions via negative log likelihood (NLL), and compared it to the NLL from a random policy. The NLL value of the velocity and reaction time policies were better than choosing randomly (two-sided t-test, velocity: p=0.0026; reaction time: p=0.028).

Overall, using vigor in probe trials as a proxy for subjective value was informative, as the results were better than chance. Reaction time was a better predictor than velocity, allowing one to predict with roughly 60% accuracy the choices made by the participants. This compares with the ceiling performance of roughly 80% accuracy when subjective value was estimated from the actual choices.

### Vigor patterns in decision trials

In the decision trials the participants expressed their choices with a saccade. We asked whether vigor patterns in the decision trials carried information about the contents of the trial.

The top plot of Fig. 5 displays time to decision (deliberation time) as well as saccade velocity as a function of trial difficulty. To quantify trial difficulty, we measured the subjective value of the chosen option minus the subjective value of the alternative option. For example, if the participant chose the sure option, trial difficulty was the subjective value of the chosen stimulus minus 0.5 times the sum of subjective values of the two stimuli in the risky option. As this difference became more positive, the decision became easier. Indeed, easier decisions coincided with reduced deliberation time (labeled DT in the top panel of Fig. 5, F(1,149)=68.3, p=7.2×10^−14^). Saccade velocity that reported the choice tended to be low in the most difficult trials (yellow region, top panel of Fig. 5), possibly indicating that reward was uncertain. This is consistent with earlier work that reported low saccade vigor in trials with increased reward uncertainty (Seideman et al., 2018).

**Fig. 5.**
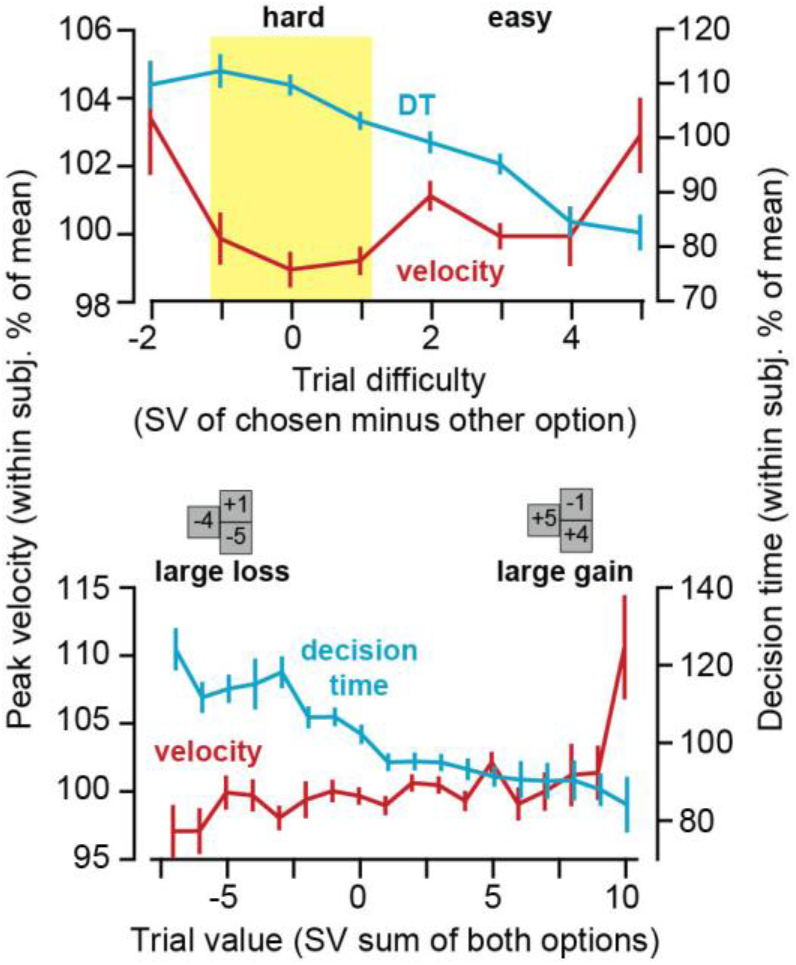
Behavior during decision trials. Decision time refers to time from trial onset to the saccade onset that indicated choice. Peak velocity refers to the velocity of the saccade that indicated choice. Top panel: trial difficulty was measured for each participant via the difference between subjective value of the chosen option minus the value of the other option. Hard choices are those in which this difference is less than or equal to 1. Easy choices are those in which this difference is greater than 1. Bottom panel: trial value was measured for each participant via the sum of the subjective values of the two options. When this sum was large and positive, both options were good, predicting gain regardless of the chosen option. When this sum was large and negative, both options were bad, thus predicting loss. Saccade velocity that indicated choice increased with trial value.

The random nature of the stimuli resulted in some trials that predicted a large loss regardless of choice, and other trials that predicted a large gain. For example, when the sure stimulus was associated with a gain, the risky option often also included stimuli that summed to a gain. Thus in this case both options were associated with gain, producing a large trial value. When we considered trial value as the sum of the subjective values of both options, a pattern emerged: when the trial predicted a loss (because both options were bad), reaction times tended to be long and saccade velocities tended to be low (lower plot of Fig. 5). As trial value increased, so did saccade vigor (velocity: F(1.275)=20.3, p=9.8×10^−6^; decision time: F(1,275)=133, p=2.1×10^−25^). The dependence of saccade vigor on trial value was present even after we normalized trials based on their difficulty (effect of sum of subjective values on decision time in easy trials: F(1,457)=130.5, p=9.3×10^−27^; in hard trials: F(1,457)=7.56; p=0.0062).

In summary, when the two options were both bad, forecasting a loss, velocity of the saccade that reported the choice was low. As the value of the trial improved, forecasting a gain, saccade velocity increased. Thus, in both probe and decision trials, saccade vigor tended to be low for trials that predicted loss, and high for stimuli that predicted gain.

## Discussion

The brain makes decisions based on subjective valuation of the available options. Yet, how we value an option is a hidden variable that cannot be measured directly. Rather, it must be inferred from our decisions. Is there a component of behavior other than choice that can serve as a proxy for subjective valuation?

Here, we presented abstract visual stimuli that participants learned to associate with gains or losses. We inferred the values that each participant assigned to the stimuli from their choices in decision trials. As expected, some participants learned a steep value function that strongly differentiated the various stimuli, whereas others learned a shallow function. In probe trials, we presented the participants a single stimulus near the fixation spot, indicating the value of that trial, and asked them to make a saccade to a peripheral target. The reaction time and peak velocity of that saccade carried information about the value of the stimulus: saccade vigor was lowest for stimuli that forecasted loss, highest for stimuli that predicted gain. Notably, saccade vigor was more strongly modulated in those participants who had also learned a steeper value function.

As expected, some participants valued a given stimulus more, whereas others valued it less. A critical question was whether between-subject differences in valuation could be gleaned from the between-subject differences in their patterns of vigor. We found that for a given stimulus (thus a constant objective value) there was a relationship between subjective value and vigor: individuals that assigned larger subjective value to a stimulus also tended to move with greater vigor in response to that stimulus.

Finally, we asked how well vigor measurements in probe trials could act as a proxy for subjective value, and thus predict choices that individuals would make in decision trials. We found that reaction time was a better predictor that peak velocity, allowing one to predict with roughly 60% accuracy the choices made by the participants (as compared to a ceiling of about 80% accuracy, as described by a model fitted to the decision trials). Thus, vigor in probe trials was modulated by subjective value, affording a modest ability to predict individual preferences during decision making.

### Estimating subjective value

Estimating subjective value usually relies on a concept called certainty equivalence (CE): if the option is a risky one that has 50% probability of producing one of two results, then the CE will be the mean of the subjective values of the two results. CE could be directly reported by the participants (Grether and Plott, 1979), or via fitting of a logistic function between the a fixed risky option and the variable sure option, or even using a psychological adaptive method such as PEST (parameter estimation by sequential testing) in which each option depends on the choice made in the previous option (Bostic et al., 1990;Christopoulos et al., 2009;Stauffer et al., 2014).

Here we employed a different approach: we implemented a simple neural network, which we found to be an efficient way to infer subjective values from the patterns of choice. Our specific learning rule relied on a loss function that guaranteed that the result would produce the optimum prediction of choices made by each participant. Our approach had the advantage that it allowed us to use a relatively small number of decision trials (600) in which all stimuli were chosen at random. The method produced reasonable results: subjective valuation correlated strongly with objective value (Fig. 1F), producing correct prediction of choice on roughly 81% of the trials.

However, we analyzed the data based on an assumption of stationarity of subject values. That is, we assumed that subjective value was constant throughout the decision trials. We provided 100 baseline trials which provided information about value of each stimulus to the participants before the decision trials began, but our assumption is clearly a simplification. Unfortunately, it is difficult to analyze the data without the assumption of stationarity because in that case one must assume a learning model, which introduces further unknown parameters that require fitting to behavior. However, if such an approach could be pursued, then one could estimate subjective value as a function of time, and look for correlations with vigor.

### Subjective value monotonically varies with saccade vigor

The main question that we wished to ask was whether vigor of a movement was a monotonic function of its subjective value across the range that spanned loss to gain. While subjective value may be lower for a loss, the stimulus that predicts a loss may gather equal or greater attention than the stimulus that predicts gain. The neural circuits that influence saccade vigor are affected by both subjective valuation (Platt and Glimcher, 1999) and attention (Leathers and Olson, 2012), making it unclear whether vigor would be influenced by one or the other.

Reaction time and velocity of a saccade are variables that are controlled by activity of neurons in the superior colliculus (Dorris and Munoz, 1995;Dorris et al., 1997;Sparks and Hu, 1999;Ratcliff et al., 2003;Smalianchuk et al., 2018). Collicular activity is in turn influenced by the excitatory inputs it receives from the cerebral cortex, and the inhibitory inputs that it receives from the basal ganglia. The cortical inputs include projections from the frontal eye field (FEF) and lateral intraparietal area (LIP), both of which house neurons that tend to respond more strongly to stimuli that predict greater reward (Glaser et al., 2016;Platt and Glimcher, 1999;Louie and Glimcher, 2010). The basal ganglia projections are from the substantia nigra reticulata (SNr), which houses inhibitory neurons that change their discharge in response to magnitude of reward (Sato and Hikosaka, 2002;Yasuda et al., 2012;Yasuda and Hikosaka, 2017). Thus, subjective valuation of a rewarding stimulus could, in principle, be reflected in an increase in the excitatory inputs to the colliculus from the cortex, and a decrease in the inhibitory inputs from the SNr, resulting in a saccade that has a shorter reaction time and greater velocity.

However, the cortical and basal ganglia inputs to the colliculus are also affected by attentional demands of the stimulus. For example, firing rates of neurons in the regions that project to the colliculus are not monotonically driven by value of the stimulus. For example, LIP neurons that respond with greater activity to more rewarding stimuli also respond more strongly to stimuli that predict a stronger loss (Leathers and Olson, 2012). In the basal ganglia, activity of SNr neurons is controlled directly and indirectly by neurons in the striatum, which in turn are modulated by dopamine. Dopamine regulates how the striatal neurons respond to cortical inputs. However, while increased dopamine release before onset of a movement tends to invigorate that movement(Kawagoe et al., 2004), some dopaminergic neurons show increased activity in response to a reward predicting stimulus, while others respond with greater activity to both reward and punishment predicting stimuli (Matsumoto and Hikosaka, 2009).

Thus, assuming that saccade vigor is a reflection of excitatory cortical and inhibitory basal ganglia inputs to the colliculus, the current neurophysiological data do not specify whether vigor should be a monotonic function of stimulus value, growing from loss to gain, or whether vigor should be a U-shaped function, showing increased activity both for large gains and large losses.

Here our results unequivocally demonstrate that saccade vigor grows monotonically with subjective value across the range that spans from loss to gain, and is not a U-shaped function. It is possible that in some of the earlier studies in which cortical and dopaminergic activity increased with punishment, the movement that followed may have been expressed with greater vigor (for example, increased rate of blinking).

Furthermore, we found that it was possible to infer some of the between-subject differences in valuation from the between-subject differences in patterns of vigor. This is reminiscent of an earlier study that found a monkey that did not show vigor sensitivity to reward also lacked dopaminergic sensitivity to stimuli that predicted reward (Kawagoe et al., 2004).

### Limitations

In our experiment the participants learned the value of the stimuli through observation (probe trials) and choice (decision trials), but we analyzed the data as if the subject values were constant throughout the decision trials. A better approach would be to have a real-time estimate of subjective values during the task. However, such an approach would require fitting behavior to a learning model, which introduces new parameters in the estimation problem. That approach remains to be developed.

In probe trials, the stimulus predicted a loss or gain if the participant performed the correct action (saccade to target). However, if the participant performed an incorrect action (or no action), the consequence was a large loss. Thus, in probe trials the participant could prevent a large loss by performing the correct action, but could not prevent the smaller loss associated the stimulus. In a different design in which the stimulus predicts a loss, but the correct action can prevent it, vigor of that action will likely grow with magnitude of loss. That is, if the correct action can aid in prevention of a loss, then we speculate that vigor would no longer exhibit the pattern we found here. This conjecture remains to be tested.

To test whether subjective value affects vigor, we relied on the fact that among participants, a given stimulus was associated with a range of subjective values. This between-subject analysis revealed that individuals who valued a stimulus more tended to also exhibit greater vigor. However, in order to conclusively infer a causal relationship between subject value and vigor we would need to test whether within participant changes in subjective value produce changes in vigor. One way with which subjective valuation may be increased is via expenditure of effort: individuals who expend effort in order to acquire a particular reward tend to increase the value that they assign that reward. With saccades, effort expenditure can be modulated via eccentricity (Yoon et al., 2018). Future work is needed to explore the within-subject changes in valuation with their vigor.

## Acknowledgements

The work was supported by grants from the NIH (5-R01-NS078311, 1-R01-NS096083), the Office of Naval Research (N00014-15-1-2312), and the National Science Foundation (CNS-1714623).

